# Porcine model elucidates function of p53 isoform in carcinogenesis

**DOI:** 10.1101/2020.09.07.286005

**Authors:** Guanglin Niu, Isabel Hellmuth, Tatiana Flisikowska, Hubert Pausch, Beate Rieblinger, Alexander Carrapeiro, Benjamin Schade, Brigitte Böhm, Eva Kappe, Konrad Fischer, Bernhard Klinger, Katja Steiger, Reiner Burgkart, Jean-Christophe Bourdon, Dieter Saur, Alexander Kind, Angelika Schnieke, Krzysztof Flisikowski

## Abstract

**Background:** The pig has long been an important animal species for biomedical research. Recent years has also seen an increasing number of genetically engineered pig models of human diseases including cancer. We previously generated pigs with a modified *TP53* allele which carries a Cre-removable transcriptional stop signal in intron 1, and an oncogenic mutation *TP53^R167H^* (orthologous to human *TP53^R175H^* and mouse *Trp53^R172H^*) in exon 5. Pigs with the unrecombined mutant allele (fl*TP53^R167H^*) develop osteosarcoma (OS) in aged heterozygous and young homozygous animals. In addition, some homozygous animals also developed nephroblastomas and lymphomas. This observation suggested that *TP53* gene dysfunction is itself the key initiator of tumorigenesis, but raises the question which aspects of the *TP53* regulation leads to the development of such a narrow tumour spectrum, mainly OS.

**Methods:** We performed a series of molecular and cellular analyses to study the regulation of *TP53* and its family members in both healthy tissue and tumours (n= 48) from fl*TP53^R167H^* pigs. Human OS cell lines were used to prove relevance to human patients.

**Results:** Molecular analyses of p53 revealed the presence of two internal *TP53* promoters (Pint and P2) equivalent to those found in human. Consequently, both pigs and human express *TP53* isoforms. Data presented here strongly suggest that P2-driven expression of the mutant R167H-Δ152p53 isoform (equivalent to the human R175H-Δ160p53 isoform) and its circular counterpart *circTP53* determine the tumour spectrum and play a critical role in the malignant transformation of bones, kidney or spleen in fl*TP53^R167H^* pigs. The detection of Δ152p53 isoform mRNA in serum is indicative of tumorigenesis. Furthermore, we showed a tissue-specific p53-dependent deregulation of the p63 and p73 isoforms in these tumours.

**Conclusions:** This study highlights important species-specific differences in the transcriptional regulation of *TP53*. For the first time a *circTP53* RNA was identified. Results indicate that the Δ152p53 isoform, its circular *circTP53* and p53 family members, TAp63δ and TAp73δ, likely play a role in the malignant transformation of bone and other tumours. Considering the similarities of *TP53* regulation between pig and human, these observations provide useful pointers for further investigation into isoform function including the novel *circTP53* in both the pig model and human patients.

## Introduction

The pig is becoming an important animal model for human cancers research, translational studies and preclinical trials (1,2). We have generated genetically engineered (GE) pigs with mutations in key tumour suppressor genes and proto-oncogenes (3), which can be used to support preclinical studies (4). As part of this program we generated pigs modelled on GE mice carrying a Cre-inducible oncogenic mutant *Trp5^R172H^* allele, in which a floxed transcriptional stop cassette is inserted into *Trp53* intron 1 to block transcription of the mutated allele (5). The latent allele can be experimentally activated by Cre recombinase to express mutant p53^R172H^ and so model spontaneous somatic *Trp53* mutation in chosen organs. These mice have been used very successfully to model a series of cancers e.g. pancreatic, breast and lung (6–9). Similarly, our pig line carries an engineered endogenous *TP53* gene with a floxed transcriptional stop signal in intron 1 and a point mutation resulting in an arginine to histidine substitution at codon 167 in exon 5 (*TP53^R167H^*, orthologous to mouse *Trp53^R172H^* and human *TP53^R175H^*) (10). As this pig line was being established it became apparent that animals with the latent non-induced allele (designated here as fl *TP53^R167H^*) in both heterozygous and homozygous form developed osteosarcomas (OS) (11) and less frequently kidney tumours and lymphomas.

This phenotype contrasts sharply with that reported for mice carrying the apparently equivalent non-induced *Trp53^R172H^* allele, which mostly develop lymphoma (12). None of the murine *Trp53* mutant strains so far generated show a preponderance of OS in the tumour spectrum (12, 13). These include Cre-inducible point mutations (13), partial deletions of the *Trp53* coding region (5), and transgenic lines that overexpress mutant *Trp53^R270H^* and *Trp53^R172H^* from exogenous promoters (14). To increase the incidence of OS, several mouse strains expressing osteoblast- and mesenchymal-specific Cre coupled with conditional *p53* and *Rb1* mutant alleles were generated (15).

In human, OS is the major form of primary bone cancer (16), which predominantly affects young people and is highly malignant, requiring aggressive surgical resection and cytotoxic chemotherapy (17). The 5-year survival rate has remained unchanged over the past 20 years, at ~60% for patients with primary osteosarcoma and ~20% for patients with metastatic disease (18). Most OS are sporadic and of unknown cause, but increased incidence is associated with Li-Fraumeni syndrome caused by germ line mutation of *TP53* (19), (20).

The role of *TP53* mutations in numerous cancers has been extensively documented (21). Yet, potentially important aspects of *TP53* gene function still remain unclear, including the role of the various p53 isoforms. Human *TP53* is known to express at least nine different mRNA transcripts (22) and at least twelve protein isoforms (23), with transcription initiated by two promoters: P1 at the 5’ end and P2 in intron 4; alternative splicing across introns 2 and 9; and alternative translation initiated at internal start codons 40, 133 and 160. The internal promoter P2 originates the Δ133p53 and Δ160p53 isoforms in humans. While the function of Δ160p53 is less studied, the Δ133p53 is involved in the regulation of replicative cellular senescence (24), angiogenesis, cytokine secretion/immune response and tumour progression in some cancer types (25).

Of note, the p53 isoforms and the internal promoters have so far been described for humans, primates, zebrafish, drosophila and mouse (23). Here we show that the porcine *TP53* locus contains internal Pint and P2 promoters equivalent to those found in human, resulting in expression of Δ152p53α isoform and its circular counterpart *circTP53*. Both are involved in the development of OS and others tumours in the fl*TP53^R167H^* pigs, and the detection of Δ152p53α isoform in serum is indicative of tumorigenesis. Tissue-specific p53-dependent deregulation of p63 and p73 isoforms could be observed in these tumours.

## Results

### Cancer spectrum in flTP53^R167H^ pigs

Since our publication describing the generation of fl*TP53^R167H^* pigs (10), a total of 29 fl *TP53^R167H/+^* heterozygous and ten fl*TP53^R167H/R167H^* homozygous pigs were examined by necropsy. All had reached sexual maturity. Animals were sacrificed as soon as they showed symptoms effecting their wellbeing or in case of the heterozygous animals latest at the age of 36 months. By this time 18 of the 29 heterozygous animals had developed tumours, all of which were classified histologically as osteoblastic OS. Spontaneous OS is very rare in wildtype pigs (11, 26). All homozygous pigs had OS by the age of 16 months or earlier, and five of these (aged 7 - 14 months) also had nephroblastoma (kidney) and diffuse large B-cell lymphomas (DLBCL, spleen). In each case the anatomical location and histological analyses of the tumours resembled those of the human juvenile cancer. As in children, OS in the homozygous pigs arise primarily around the knee joints, femur and tibia, all regions of active bone growth.

### Identification of a Δ152p53α isoform in pigs

The observation that the tumour repertoire was restricted to a few tissue types (100% OS, 22% nephroblastoma and 17% B-cell lymphomas) lead to the question, what aspects of *TP53* expression or absents thereof supports the formation of mainly OS rather than other tumour entities? To answer this question we first establish if expression of the floxed fl*TP53^R167H^* allele was silenced in all organs.

A series of RT-PCR primers hybridising to different porcine *TP53* exons (Figure 1a) were used to screen RNA samples from various organs (n= 11) of fl*TP53^R167H/R167H^* homozygous pigs (n= 3). No mRNA expression was detected using RT-PCR primers specific for exon 1 to exon 2 (1F/1R), or exon 2 to 4 (2F/2R) in all tissues analysed including bone and OS samples (Figure 1b), confirming that the LSL efficiently blocked transcription. However, RT-PCR using primers between exons 5 to 11 (5F/9R, see Figure 1a) revealed a tissue-specific expression (Figure 1c), where the strongest RT-PCR signal was obtained for bone, lymph node, and kidney tissues. This indicated the presence of a promoter element in intron 4 similar to human. No expression of the short *TP53* mRNA was detected in heart and aorta. To determine the start and termination of the porcine *TP53* isoform 5’ and 3’ rapid amplification of cDNA end (RACE) analysis was performed (Figure 1d and e). This placed the start of the isoform 11 base pairs upstream of exon 5 with a possible ATG at amino acid position 152 (Δ152p53 isoform), which translates in frame and terminates at the same stop codon as the full length p53 and corresponds to *TP53α* isoform (Supplemental Figure 1). The identified Δ152p53α mRNA has a length of 1295 bp, the nucleotide sequence was verified by sequencing.

**Figure 1.**
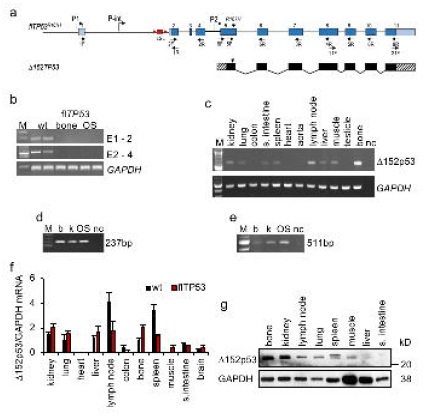
Identification of Δ152p53α isoform in pigs. **(a)** Schematic representation of the *TP53* gene and the Δ152p53α mRNA. **(b)** Expression analysis across exon 1 to 2 (186bp), exon 2 to 4 (591bp) of bone samples from wild-type pigs and healthy bone and OS samples from *flTP53^R167H/R167H^* pigs showing lack of expression for mutant TP53. **(c)** Expression analysis across exon 5 to 11 (492bp) for 11 different organ samples from *flTP53^R167H/R167H^* pigs. **(d)** 5’ RACE using a reverse primer hybridising to exon 6 of *TP53* and resulting in a 237bp fragment and **(e)** 3’ RACE using forward primer hybridising to exon 11 of *TP53* resulting in a 511bp fragment. Healthy bone (b), kidney (k) and osteosarcoma (OS) samples from *flTP53^R167H/R167H^* pigs,. M – marker, nc – negative control. **(f)** Quantitative RT-PCR analysis using primers hybridising to extended exon 5 and exon 6 and detecting the Δ152p53 isoform mRNA expression in different tissues of *flTP5^R167H/R167H^* (n= 6) and wild type (n= 3) pigs*. GAPDH* was used as a reference gene to calculate the relative expression levels. **(g)** Detection of Δ152p53 isoform by western blot analysis using Sapu antibody in different tissues from *flTP5^R167H/R167H^* pigs.

### Expression of Δ152p53α isoform is highest in organs susceptible to cancer development

By using primers (9F/6R) specific for the Δ152p53α isoform its expression was quantified by qPCR in different healthy tissues (n= 11) from fl*TP53^R167H^* homozygous (n= 10) and wild type (n= 3) pigs. The highest Δ152p53α mRNA expression in fl*TP53^R167H^* homozygous pigs was observed in lymph nodes, spleen, kidney and bone (Figure 1f), all organs prone to tumour development in mutant pigs. Except for lymph node, colon and small intestine the Δ152p53α mRNA expression was generally lower in wild type samples, than in fl*TP53^R167H^* homozygous samples.

### The porcine Δ152p53α isoform encodes a 30kDa protein

The methionine codon 152 in pigs corresponds to codon 160 in human p53, which is located within a highly conserved Kozak sequence (27). An *in silico* analysis predicted that the 1295 bp mRNA encodes a 224 amino acid N-terminal truncated Δ152p53α isoform using the same reading frame as full length p53. The predicted porcine isoform shares 88% amino acid sequence homology with the human Δ160p53α isoform, which uses the Δ133/160p53 alternative transcriptional initiation site in exon 5 (27) and contains part of the highly conserved p53 DNA binding region (23). To determine whether the porcine *TP53* alternative transcript can produce Δ152p53α protein, western blot analysis of different healthy tissue samples from fl*TP53^R167H/R167H^* homozygous pigs was performed using the polyclonal sheep SAPU antibody recognising all human p53 isoforms. Two bands (doublet) of ~30kD were detected (Figure 1g). As with the mRNA highest protein expression was observed in bone, kidney, lymph node, and spleen. The size of the western blot bands was comparable to the human Δ160p53 isoforms (27), and most likely represent splice variant or post-translational modifications of R167H-Δ152p53 proteins.

### Identification of internal promoters in the porcine TP53 gene

Results above clearly indicated the presence of at least one internal promoter. To determine the location of all possible internal promoters a nucleotide sequence alignment between the porcine (NC_010454, GenBank Sscrofa 11.1) and human (NC_000017, Genbank GRCh38.p12) *TP53* gene was carried out. Apart of exons, three genomic regions with high homology (> 70% similarity) were detected (−1877bp to +11bp, +2125bp to +3200bp and +9985bp to +10470bp; relative to the major transcription start site, mTSS). These porcine *TP53* regions correspond to the locations of the P1 (5’end), Pint (intron 1) and P2 (intron 4) promoters in human *TP53*. As for the human sequence, the porcine intron 4 sequence (P2) contains binding sites for the transcription factors NFIC, Hltf and SPI1. In contrast, the sequence of mouse *Trp53* intron 4 showed less than 50% homology to human *TP53*, and lacks binding sites for these transcription factors (Supplemental Figure 2). Interestingly the NFIC transcription factor regulates osteoblast differentiation (28), and can promote or suppress the development of various cancers (29) via epigenetic changes (30). The SPI1 transcription factor regulates alternative transcription of target genes, and both Hltf and SPI1 are frequently mutated in paediatric cancers (31,32).

To confirm promoter function five luciferase reporter constructs were generated containing the P1 promoter (−2000bp to TSS), the putative P2 promoter (intron 4) and three different fragments covering the first intron (Pint_1, Pint_2, Pint_3), see Figure 2a. Luciferase expression under the control of the SV40 promoter was used as positive control.

**Figure 2.**
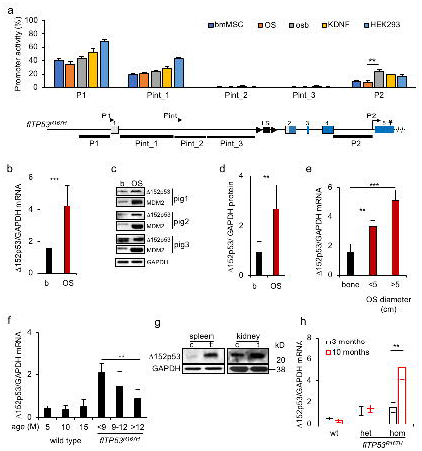
Identification of porcine *TP53* promoters and expression of Δ152p53α isoform in osteosarcomas. **(a)** Dual-luciferase assay: Luciferase was expressed from the SV40 promoter and used to normalise the Renilla expression under the control of the putative promoter fragments. Their location relative to the gene structure is depicted. Values represent mean ± standard deviation, six transfections per construct. Promoterless luciferase vector was used as a negative control. bmMSC – bone marrow mesenchymal stem cells, osb – porcine fl*TP53^R167H/R167H^* osteoblasts, OS – porcine *flTP53^R167H/R167H^* osteosarcoma cells, KDNF – porcine kidney fibroblasts, HEK293 – human embryonic kidney cell line. **(b)** Quantitative PCR results of Δ152p53α mRNA expression in OS (n = 48) and matched healthy bone samples. **(c)** Representative western blots showing Δ152p53α protein and MDM2 expression in OS and healthy matched bone samples of homozygous *flTP53^R167H^* pigs. **(d)** Quantitative measurements of proteins in OS (n=10) and healthy matched bone samples from homozygous *flTP53^R167H^* pigs. **(e)** The Δ152p53α mRNA expression in small (– 5 cm, n= 29) versus large (> 5 cm, n= 19) tumours. **(f)** Age-dependent Δ152p53α mRNA expression in healthy bones of homozygous fl*TP53^R167H^* (n= 10) and wild type (n= 3) pigs. **(g)** Western blots showing Δ152p53α isoform expression kidney and spleen tumours and healthy matched tissues of *flTP5^R167H/R167H^* pigs. **(h)** QPCR analysis of Δ152p53α mRNA expression in blood exosomes from fl*TP53^R167H^* heterozygous (n= 6), homozygous (n= 6) and wild type (n= 3) pigs aged 3 and 10 months.

Luciferase activity was measured after transfection of various porcine primary cells: wild type bone marrow mesenchymal stem cells (bmMSC) and kidney fibroblasts; fl*TP53^R167H/R167H^* OS cells and healthy osteoblasts; and human HEK293 cells. In comparison to the positive control two of the putative promoter fragments led to moderate (P1, 40-70% and Pint_1, 30-45%) and one to low (P2, 10-25%) luciferase activity (Figure 2a). These three promoter fragments shared the greatest homology with the corresponding human *TP53* sequence. The *in silico* and experimental data provide evidence for the presence of internal promoters within intron 1 (P-int) and intron 4 (P2) of the porcine *TP53*, similar to those in human *TP53*.

Expression from the two main promoters P1 and Pint is blocked by lox-stop-lox cassette in fl*TP53^R167H^* homozygous pigs. Additional deletion of the P2 promoter fragment should result in a complete inactivation of porcine *TP53*. Using the Crispr-Cas9 system and two guide RNAs a DNA fragment from 9665 to 10374bp of pig *TP53* (NC_010454) was excised in OS cells from fl*TP53^R167H^* homozygous pigs (Supplemental figure 3a). RT-PCR analysis showed no *TP53* expression in the edited porcine OS cells (Supplemental figure 3b), confirming that the P2 promoter is responsible for the Δ152p53α expression.

### The R167H-Δ152p53αisoform is overexpressed in tumours

To assess the RNA and protein expression of Δ152p53α isoform in cancer, we analysed tumours collected from fl*TP53^R167H^* pigs. Quantitative PCR and western blot revealed more than three-fold (*P* < 0.001) higher expression of R167H-Δ152p53α isoform in OS compared to matched healthy bones (Figure 2b-d). The level of Δ152p53α mRNA expression increased (*P* = 1.17 x 10^-5^) with OS tumour size (Figure 2e).

The expression level of the Δ152p53α mRNA in healthy bones correlated with the onset of symptoms in homozygous fl*TP53^R167H^* pigs (Figure 2f). The higher R167H-Δ152p53α was expressed the earlier tumours became evident. In wild type pigs the level of Δ152p53α expression remained unchanged with age (Figure 2f).

The above results hinted that overexpression of the mutant R167H-Δ152p53α isoform was essential for tumorigenesis. To assess if this also applied to other tumour types samples from the nephroblastomas and B-cell lymphomas were analysed. As for the OS, an overexpression of Δ152p53α isoform was observed in kidney and spleen tumours (Figure 2g). Finally, we also tested if the orthologous Δ133/160p53 mRNA was expressed in human OS samples. These too were positive (Supplemental figure 4).

Overexpression of the Δ152p53α isoform seemed essential for tumour growth. To confirm this, and to assess if the effect was due to the expression of the wildtype or mutant isoform, a proliferation assay was performed. Pig OS cells transfected with an expression vector carrying the wild type Δ152p53α or mutant R167H Δ152p53α cDNA sequence under the control of the CAG promoter showed significantly increased proliferation compared to those transfected with control GFP vector (Supplemental figure 5). The highest proliferative increase was obtained for the mutant isoform.

The MDM2 oncoprotein is a key regulator of p53 expression, which is stabilized by mutant p53 (33). To determine if the mutated Δ152p53α protein isoform retains this function, western blot analysis was carried out and showed an increase in MDM2 protein in OS tumour samples (Figure 2c). Taken together, these data indicate that the mutated Δ152p53α isoform plays a critical role in the malignant transformation of bones, kidney or spleen in fl*TP53^R167H^* pigs.

### Blood exosomal Δ152p53αexpression is indicative of tumorigenesis

In order to investigate whether the increased expression of R167H-Δ152p53α mRNA during malignancy can be detected in serum samples and used as a biomarker, exosomes were isolated from *flTP53^R167H^* heterozygous (n= 6), homozygous (n= 6) and wild type (n= 3) pigs aged 3 and 10 months. At the age of 3 month all animals were disease free while at the age of 10 months the *flTP53^R167H^* homozygous pigs showed first signs of cancer, confirmed later by necropsy. QPCR showed a 6-fold higher level (P< 0.01) of R167H-Δ152p53α mRNA in exosomes from 10-months old *flTP53^R167H^* homozygous pigs than from the same pigs at the age of 3-months (Figure 2h). The exosomal Δ152p53 mRNA expression in *flTP53^R167H^* heterozygous and wild type pigs was low and unchanged over the same time period (Figure 2h). This data suggests that detection of Δ152p53α isoform mRNA in serum is indicative of tumorigenesis.

### The DNA methylation in the P2 promoter is negatively correlated with the mutant-Δ152p53αexpression in OS cells and tumours

The P2 promoter showed an increased activity in tumours. It has been reported that intragenic *TP53* methylation differs between normal and transformed human colorectal cancer cell lines (34). To evaluate if epigenetic changes correlate with altered expression levels a comparison of the DNA methylation in healthy osteoblasts and OS cells derived from *flTP53^R167H/R167H^* pigs was carried out. CpG regions in the P1 promoter, intron 1 (fragments Pint_1, Pint_2, Pint_3), P2 promoter and exon 5 were identified. DNA methylation at four or more CpG sites was analysed for each region. Overall, CpG methylation was higher in the gene body than in the P1 promoter (Figure 3a), which is consistent with *TP53* methylation pattern in human (35). The highest DNA methylation (> 80%) was observed in Pint_2 and Pint_3 fragments that showed no promoter activity. The promoter regions of P1, Pint_1 and P2 showed 18%, 28%, and 64% DNA methylation in healthy osteoblasts, respectively. The DNA methylation at five CpG sites in Pint_1, P2 and exon 5 was significantly lower in OS than in healthy osteoblasts cells (Figure 3a). Specifically the following CpG sites showed lower DNA methylation in OS cells: in Pint_1 - CpG3 (−19%, *P* < 0.01), in the P2 promoter - CpG2 (−33%, *P* < 0.001) and CpG3 (−28%, *P* < 0.001), and within the Kozak sequence - CpG5 (−37%, *P* < 0.001) and CpG6 (−24%, *P* < 0.01). The level of DNA methylation in osteoblasts and OS cells inversely correlated with promoter activity (Figure 3b).

**Figure 3.**
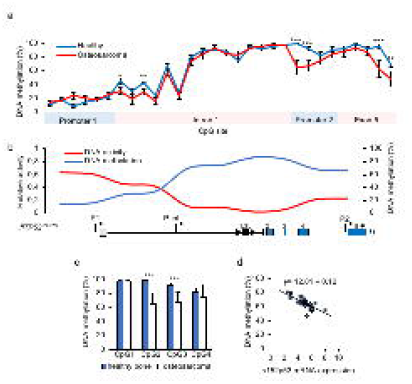
DNA methylation level of porcine *TP53* promoters. **(a)** DNA methylation analysis of CpGs located in putative promoter regions *in TP53* in osteoblasts and OS cells of *flTP53^R167H/R167H^* pigs. **(b)** The correlation plot for promoter activity and DNA methylation. The values on the plot are aligned to the *TP53* gene structure. **(c)** Pyrosequencing analysis of four CpG sites in P2 promoter (P2 fragment) in OS (n = 48) and matched healthy bone (n = 29) of fl*TP53^R167H^* pigs. Values shown represent mean ± SD. ** *P* < 0.01, *** *P* < 0.001. **(d)** Regression analysis between CpG methylation at site 2 in P2 promoter and Δ152p53α mRNA expression in OS (y= 12.81 −0.12, *P*= 2.9 x 10^-7^).

The same analysis was carried out for OS tissue samples. Compared to healthy bone the same five CpG sites as mentioned above showed significantly reduced DNA methylation in OS. The level of DNA methylation was as follows: in Pint_1 - CpG3 (41% vs. 22%, healthy bone vs. OS, *P* < 0.01), in the P2 promoter region - CpG2 (99% vs. 62%, *P* < 0.0001) and CpG3 (96% vs. 65%, *P* < 0.0001) site, and CpG5 (94% vs 63%, *P* < 0.01) and CpG6 (67% vs 52%, *P* < 0.01) site within the Kozak sequence (Figure 3c, Supplemental Figure 6e). CpGs in this region have been shown to be differentially methylated for the Δ133/160 isoforms in human (34).The regression analysis revealed that the decreased methylation at CpG2 and CpG3 sites in P2 promoter is correlated with higher R167H-Δ152p53α mRNA expression in pig OS (*P*= 2.9 x 10^-7^; Figure 3d).

### TP53 circular RNA enhances proliferation of OS cells

There is increasing evidence that circular RNAs (circRNAs) play a role in human cancer (36, 37), but so far no data has been reported showing expression of a circRNA for *TP53 (circTP53)*. To identify *circTP53* in *fl TP53^R167H/R167H^*, several pairs of divergent primers were designed for RT-PCR analysis resulting in the detection of four different circ*TP53* in fl*TP53^R167H/R167H^* tissues (*circTP53-1 to −4*, Figure 4a), of which *circTP53-3 was highly expressed in bone*. All were expressed from the P2 promoter and encoded within the Δ152p53α isoform, which was confirmed by Sanger sequencing and Rnase R digestion (Figure 4b). RT-PCR analysis revealed a tissue-specific expression of *circTP53 variants*, with their highest expression in bone, kidney and colon (Figure 4c). No expression of *circTP53* was found in heart and aorta, which is consistent with the lack of Δ152p53α expression in these organs. Compared to healthy bones, the *circTP53* expression was significantly increased (*P* < 0.001) in OS (Figure 4d,e).

**Figure 4.**
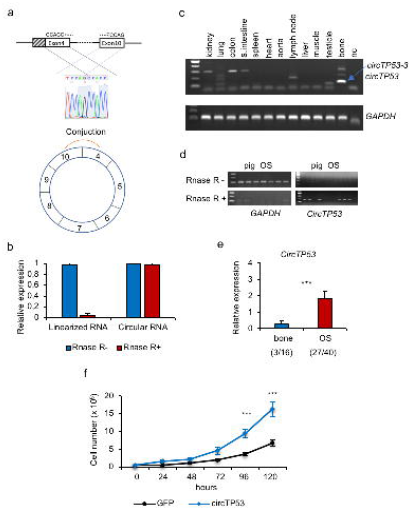
Identification of *circTP53* in *flTP53^R167H^* pigs. **(a)** Schematic presentation of *circTP53* identification and confirmation by sanger sequencing. **(b)** Q-RT-PCR results showing the effect of RNAase R digestion. **(c)** RT-PCR analysis of circTP53 in different tissues in *flTP5^R167H/R167H^* pigs. **(d)** RT-PCR amplification showing the effect of Rnase R digestion on *circTP53* and *GAPDH* amplification. **(e)** Relative expression of *circTP53* in healthy bones and OS. In parenthesis the number of *circTP53* positive to analysed samples is shown. **(f)** Proliferation assay in pig fl*TP53^R167H/R167H^* OS cells transfected with *circTP53* overexpression vector, consisting of exon 5 to exon 9 (circ*TP53-3*) under the control of CMV promoter.

To test, whether the newly detected *circTP53* has any effect on cell proliferation, an overexpression vector, consisting of exon 5 to exon 9 (circ*TP53-3*) and the R167H mutation under the control of CMV promoter, was generated. As shown in Figure 4f, the proliferation of pig OS cells transfected with *circTP53* overexpression vector was significantly increased (*P* < 0.001) compared to cells transfected with a control GFP vector.

### Altered expression of p63 and p73 in tumours from *flTP53^R167H^* pigs

The p53 family includes two other members: p63 and p73, all three genes are structurally similar, and have been implicated in cell regulation and cancer (38, 39). This raised the question whether the *TP53* mutation also influenced the expression of p63 and p73 in fl *TP53^R167H^* pigs.

#### p63

has two major isoforms, TAp63 and ΔNp63, which have different roles in tumorigenesis. While ΔNp63 isoform promotes, the TAp63 suppresses tumour growth in mice (40). Different tissues (n= 11) from fl*TP53^R167H^* homozygous (n= 10) and wild type (n= 3) pigs were analysed by qPCR with primers located in exon 3 and exon 4. The expression of the detected *TP63* mRNA variant was significantly lower in heart, lung, bones and higher in lymph node, colon, spleen in fl*TP53^R167H^* homozygous compared to wild type pig tissues (Figure 5a). Western blot analysis revealed tissue specific expression of two TAp63 isoforms, TAp63α and TAp63δ and no expression of ΔNp63 (Figure 5b). While, the TAp63α isoform was highly expressed in wild type kidney, the TAp63δ showed an overexpression in wild type bones, and kidney and spleen tumours (Figure 5c), and variable expression in OS (Figure 5d) from fl*TP53^R167H^* pigs, indicating that the *TP53* mutation has a tissue- and tumour-specific effect on the TAp63δ isoform expression.

**Figure 5.**
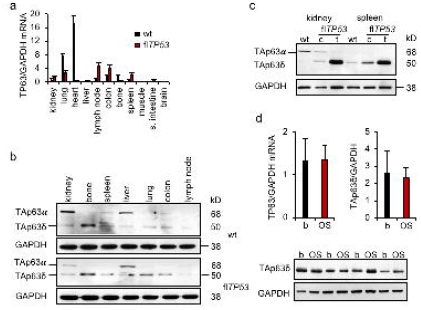
Expression of p63 in *flTP53^R167H^* pigs. **(a)** Quantitative RT-PCR analysis of different tissues (n= 11) from *flTP5^R167H/R167H^* (n= 6) and wild type (n= 3) pigs. **(b)** Representative western blots showing p63 protein expression in different tissues of *flTP5^R167H/R167H^* and wild type pigs. **(c)** Western blots showing p63 isoforms expression in healthy and tumour tissues of kidney and spleen of *flTP5^R167H/R167H^* and wild type pigs. **(d)** Quantitative PCR results of p63 mRNA expression in OS (n = 48) and matched healthy bone samples. **(e)** Representative western blots showing p63 protein expression in OS and healthy matched bone samples of homozygous *flTP53^R167H^* pigs. Quantitative measurements of protein in OS (n=10) and matched healthy bone samples of homozygous *fl TP53^R167H^* pigs.

#### p73

In humans, p73 has two different promoters, which regulate several isoforms: full length TAp73 and N-terminal truncated ΔNp73, which can be distinguished by their transactivation functions (41, 42). It has been suggested that TAp73 has a tumour suppressor function similar to that of p53, whereas ΔNp73 isoforms would promote cell growth by regulating activities of p53 family members (43). QPCR analysis revealed a high *TP73* mRNA expression in kidney, lung, liver, colon, spleen, bone, lymph node and no expression in muscle and brain of wild type animals (Figure 6a). It was significantly lower in colon and small intestine but higher in lung, bone, spleen of fl*TP53^R167H^* homozygous pigs compared to wild type (Figure 6a). Western blot detected the TAp73α isoform in kidney and liver and the TAp73δ isoforms in spleen, liver, lung of fl*TP53^R167H^* homozygous and wild type pigs (Figure 6b). Compared to wild-type, an overexpression of TAp73δ isoform was observed in healthy bones (Figure 6b) and tumours (Figure 6c, d) from fl*TP53^R167H^* homozygous pigs.

**Figure 6.**
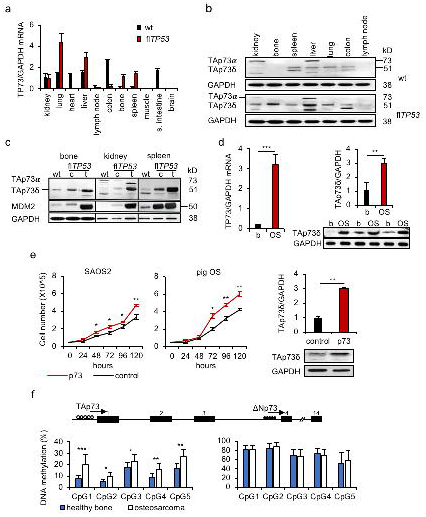
Expression of p73 in *flTP53^R167H^* pigs. **(a)** Quantitative RT-PCR analysis of p73 mRNA expression in different tissues (n= 11) of *flTP5^R167H/R167H^* (n= 6) and wild type (n= 3) pigs. **(b)** Western blots showing the tissue specific p73 isoforms expression in *flTP5^R167H/R167H^* and wild type pigs. **(c)** Western blots showing p73 isoforms and MDM2 proteins expression in bone, kidney, and spleen tumours and healthy matched tissues of *flTP5^R167H/R167H^* and wild type pigs. **(d)** Quantitative PCR (left) and western blot (right) measurements showing p73 expression in OS (n = 48) and matched healthy bone samples. Representative western blots (bottom) showing TAp73δ isoform expression in OS and matched healthy bone samples of homozygous *flTP53^R167H^* pigs. Quantitative measurement of western blots was performed for OS (n=10) and matched healthy bone (n= 10) samples of homozygous *flTP53^R167H^* pigs. **(e)** Proliferation assay in human SAOS2 (left) and pig *flTP53^R167H/R167H^* OS (middle) cells. Western blot (right) showing the upregulation of the TAp73δ isoform in pig OS cells transfected with a full length of *TP73* cDNA vector. The bar plot above shows the quantitative protein measurements (n= 3) of western blots. **(f)** DNA methylation analysis of P1 and P2 promotor regions of *TP73* in OS (n = 48) and matched healthy bone samples of *fl TP53^R157H^* pigs. Open and filled circles on the *TP73* gene structure indicate unmethylated and methylated DNA regions.

We next determined if the p73 isoforms expression is correlated with DNA methylation in the two promoter regions. A hypermethylation of the ΔNp73 promoter and a hypomethylation of the TAp73 promoter was observed. The DNA methylation at all five CpG sites of TAp73 promoter differed significantly, in particular at CpG1 (21% vs 9%, P< 0.001) between OS and matched healthy bone samples (Figure 6f), which supported the observed expression pattern of the TAp73 isoform.

To further investigate the function of p73 in tumorigenesis, a porcine *TP73* full length cDNA expression vector, containing both the TAp73 and △Np73 translation start codons, was transfected into porcine OS and human OS (SAOS2) cells. The *TP73* transfected cells showed an overexpression of the TAp73δ, no expression of △Np73 isoform, (Figure 6e) and significantly (P< 0.01) increased growth rate compared to GFP transfected control cells. These data indicated that the TAp73δ isoform is predominantly translated from the *TP73* cDNA and has an effect on the proliferation of these cells.

In summary, these analyses showed for the first time the expression of p63 and p73 isoforms in porcine tissues and the upregulation of the two isoforms (TAp63 and TAp73) presumed to have tumour suppressor function in porcine tumours. However, overexpression of TAp73δ did not lead to a reduction but to an increase in cell proliferation, which may confirm that different C-terminal variants may lack growth suppressing function.

## Discussion

It is becoming evident that p53 isoforms, including those derived from the internal P2 promoter, have implications in human cancers (23). Further elucidation of their roles, interactions, regulation and patterns of expression could open novel approaches for the prognosis and treatment of cancer. But the study of p53 isoforms and its involvement in tumorigenesis had been hampered by the fact that the main experimental mammal, the mouse, lacks internal P2 promoter activity (23). This motivated the generation of mice which ubiquitously express a Δ133p53-like protein (Δ122p53) through deletion of exon 3 and 4 (44, 45). Homozygous Δ122p53 mice show an enhanced proinflammatory phenotype and are prone to develop B-cell tumours (46) with low incidence of OS (17%) (45). Although this mouse model does not replicate the situation in human, its strongly supports the notion that p53 isoforms play a role in cancer.

The study presented here shows that the porcine *TP53* -unlike the mouse- has two internal promoters, Pint in intron 1 and P2 in intron 4. In humans the expression from the P2 promoter results in two isoforms Δ160p53 and Δ133p53. The porcine Δ152p53 protein isoform is equivalent to human Δ160p53. The pig lacks the transcription initiation site, which in humans enables translation of the Δ133p53 protein isoform. Three N-terminal variants have been observed in humans (p53α,β,χ). The porcine mRNA isolated by RT-PCR represents the Δ152p53α mRNA. However, western blot analysis indicated that N-terminal variants are present in the pig.

To our knowledge this is the first time that an association between P2 promoter activity and epigenetic modifications has been shown for normal and tumour tissue. An important finding as epigenetic modulation has been suggested as a means of restoring wild type p53 function, or inactivating mutant p53 activity in human cancer (47).

Comparison with human data show that the porcine Δ152p53α mRNA is expressed in a similar tissue-specific manner, including high expression e.g. in bone and lack of expression in heart tissue (22). Our study strongly suggests that the predominance of OS followed by nephroblastomas and B-cell lymphomas in pigs carrying the *floxed TP53^R167H^* allele is related to the higher level of Δ152p53α expression in these organs. This is further supported by the finding that pigs with early onset of OS have also higher expression of the Δ152p53α isoform in healthy bone tissue.

Expression of Δ152p53α isoform increases during tumorigenesis, which is consistent with the expression of p53 isoforms in human cancers (48). The overexpression of Δ133/Δ160p53 variants and their potential oncogenic function have been reported in lung, colon, breast and ovarian cancers and in melanoma (49–52). In the pig overexpression of the wild type and R167H mutant Δ152p53α isoform enhanced cell proliferation, finding consistent with data for mutant Δ160p53 isoform in human cancer cells (53). It was more pronounced for mutant isoforms, which would imply that the tissue specific expression of the mutant Δ152p53α isoform drives tumorigenesis in our pig model and is an indicative blood biomarker.

*TP53^R167H^* Yucatan minipig model generated by Sieren et al. (54), develops a similar tumour spectrum, but the role of the Δ152p53 in this model is unknown. It is questionable if the P2 promoter is functional in this model due to the insertion of a selectable marker gene in intron 4.

It has been suggested that one of the key mechanisms for p53 gain of function mutations is its interaction with p63/p73 (55, 56), and that the ratio of TA/ΔN p63/p73 isoforms determine their effect on tumorigenesis (57, 58). We observed an upregulation of the TAp63δ and TAp73δ isoforms in all studied tumours, and proved that experimental overexpression of the TAp73δ isoform increased proliferation of human and porcine OS cells. Further research is required to prove a direct interaction between the specific isoforms.

We present the first description of *circTP53* RNAs expressed from the P2 promoter. As with the parental Δ152p53α isoform it is upregulated in tumours such as OS. Importantly the functional study demonstrated that high *circTP53* expression increases cellular proliferation of OS cells. Similar mechanisms for gene overexpression and upregulation of circRNA was described for other genes, for example androgen receptor (AR) in prostate cancer (59). The mechanism how *circTP53* effects cell proliferation still needs to be elucidated.

## Conclusion

The biology of p53 has been studied extensively for four decades and still novel insights are being gained. Cross species comparison might help to understand the tissue specific regulation of *TP53*. Our study has highlighted the value of using the pig model. We show that the Δ152p53 isoform, its circular counterpart and the p53 family members, TAp63δ and TAp73δ, likely play a role in the malignant transformation of bone and other tumours. Considering the similarities of *TP53* regulation between pig and human, the observations we present here provide possibly useful pointers for further investigation into human *TP53*.

## Material and Methods

### Animals

Three wild type, and ten (8 males and 2 females) *flTP53^LSLR167H/R167H^* and 24 (13 males and 11 females) *flTP53^LSLR167H/+^* pigs aged 7 – 32 months, were produced by normal breeding and raised in our animal husbandry facilities with food and water provided *ad libitum*. All animal experiments were approved by the Government of upper Bavaria (permit number 55.2-1-54-2532-6-13) and performed according to the German Animal Welfare Act and European Union Normative for Care and Use of Experimental Animals.

### Necropsy examination and tumour analysis

Pigs were humanely killed and examined by complete necropsy. In total three bone samples from wild type and 48 OS samples with matched healthy bones from hetero- and homozygous fl*TP53^LSLR167H^* pigs were analysed. The size criteria for bone tumours are according to the AJCC Cancer Staging Manual (60). For histopathology analysis, organ specimens were fixed in 4% buffered formaldehyde, embedded in paraffin, sectioned (4 μm) and stained with haematoxilin and eosin (H&E). Bone specimens were first decalcified in Ossa Fixona (Waldeck GmbH, Germany). For cryosections normal and tumour samples were frozen in 2-methylbutane (OCT), and snap frozen for molecular analyses and stored at −80 °C.

### Exosomes isolation from blood

40 ml of EDTA blood was collected from *flTP53^R167H^* heterozygous (n= 6) and homozygous (n= 6) and wild type (n= 3) pigs aged 3 and 10 months. For the isolation and purification of exosomes the Exo Easy Maxi Kit (Qiagen) was used according to manufacturer’s protocol. The exosomal RNA was isolated using the RNeasy Midi Kit (Qiagen).

### Porcine primary cells

Porcine bone marrow MSCs (bmMSC) and kidney fibroblasts (KDNF) were derived in-house and cultured by standard procedures (10). To minimise phenotype changes, cells were kept as frozen stocks and cultured for 4 weeks maximum.

### Porcine osteosarcoma cell culture

Cells were derived from a minced piece of tumour digested in DMEM supplemented with 200U/ml collagenase type IV (Worthington, USA) at 37 °C for 24 h, then centrifuged at 100 r.c.f. for 5 min. Cells were resuspended in standard porcine MSC medium (10), cultured at 37°C, 5% CO_2_.

### Primers

The authors will provide all primers sequences used in this study on request.

### Reverse transcription PCR

100ng total RNA was used for cDNA synthesis using the Superscript IV (Thermo Fisher) according to manufacturer’s protocol. RT-PCR was carried out using PyroMark PCR mix (Qiagen) using primers that specifically detect the alternative smaller p53 RNA species. The RT-PCR products were verified as p53 RNA by sequencing.

### Quantitative real-time RT-PCR

QPCR was carried out using Kapa SYBR @ Fast Mix (Kapa Biosystems Pty, South Africa) and run on ABI 7500 PCR System (Applied Biosystems, USA) with default thermal cycling parameters. Reactions were performed in 10 μl volume. Samples were assayed in triplicate, relative expression was normalised to *GAPDH* expression and fold-differences were calculated by the δδCT method and statistically compared using Students t-test.

### Rnαse R digestion

RNase R treatment was carried out for 20min at 37°C using 2U RNase R (Epicenter) per 1μg of RNA. Treated RNA was directly reverse transcribed using Superscript IV (Thermo Fisher) with random decamer primers according to the manufacturer’s instructions

### 5’ and 3’ Rapid amplification of cDNA ends (RACE)

1 μg total RNA from healthy and OS samples was used for 5’ and 3’ RACE reactions with the FirstChoice RLM-RACE kit (Ambion) according to manufacturer’s protocol. Modified RNA was reverse transcribed using SuperScript IV (Thermo Fisher). The resulting cDNA was used for nested PCR.

### Pyrosequencing

Pyrosequencing assays were designed using PyroMark Assay Design 2.0 software (Qiagen). 500 ng genomic DNA was bisulphite-converted with the EZ DNA Methylation-Direct kit (Zymo Research, Irvine, USA) according to the manufacturer’s instructions. PCR samples were amplified using PyroMark PCR kit (Qiagen). PCR products were sequenced using PyroMark Q48 Advanced CpG reagents on a PyroMark Q48 Autoprep instrument (Qiagen). For assay optimalisation, a methylated (100%), non-methylated (0%) and a scale of control samples with the following DNA methylation: 25%, 50% and 75% were used. The methylated control was prepared using the CpG methyltransferase enzyme (M.SssI; Thermo Scientific). A nonmethylated control was prepared using REPLI-g Mini kit (Qiagen).

### Luciferase reporter assay

Porcine bone-marrow mesenchymal stem cells and kidney fibroblasts were cultured in DMEM supplemented with 20% FCS, 1% sodium pyruvate, 1% NEAA and 1% glutamine. Porcine osteoblasts and OS cells were cultured in DMEM/F12 supplemented with 10% FCS and 1% glutamine. HEK293 cells were cultured in DMEM supplemented with 10% FCS. All cells were plated into 24-well plates and then transfected with P1, Pint_1, Pint_2, Pint_3 and P2 psiCHECK2 plasmids (0.5 μg per well) using Lipofectamine 2000 transfection reagent (Thermo Fischer). Empty vector and SV40 promoter were used as negative and positive controls. Firefly and *Renilla* luciferase activities were measured using a *Firefly* & *Renilla* Luciferase Single Tube Assay kit (Biotium) on FLUOstar Omega (BMG Labtech). All assays were performed in triplicate.

### Porcine Δ152p53^WT^ and Δ152p53^R167H^ mRNA expression vectors and stable transfection

Both vectors comprised of the CAG promoter directing expression of *Δ152p53^WT^* or *Δ152p53^R167H^* mRNAs linked to bovine growth hormone polyA. These together with hygromycin selectable marker vector were than used for co-transfection of fl*TP53^167H/167H^* OS and wild type kidney fibroblasts porcine cells.

### Proliferation assay

Porcine OS cells were transfected with *Δ152p53^WT^, Δ152p53™^6^™*, and *circTP53* overexpression vectors by electroporation using the EMC830 electroporation system (BTX). GFP vector was used as a control. The transfected cells were selected by using 200ng/ul of hygromycin. After selection, 5 x 10^5^ cells were plated on 6 well-plates (3 times for each assay), and cells were counted after 24h, 48h, 72h, 96h and 120h of incubation using automated cell counter (Invitrogen).

### Western blot analysis

Protein was isolated using NP40 buffer, 40mg of protein was separated on a 12% SDS PAGE gel, transferred to PVDF membrane and processed using the iBind Western System (Thermo Scientific) according to the manufacturer protocol. Pig Δ152p53 isoform was detected using sheep Sapu antibody (diluted 1:1000) and horseradish peroxidase (HRP) labelled anti-sheep s36-62DD (diluted 1:2000). Pig p63, p73 and MDM2 were detected using rabbit anti-p63 monoclonal antibody ab124762 (diluted 1:1000), rabbit anti-p73 polyclonal antibody PA5-80175 (diluted 1:1000), rabbit anti-MDM2 polyclonal antibody ab260074 (diluted 1:1000) and horseradish peroxidase (HRP) labelled anti-rabbit sc-2004 (diluted 1:2000), respectively. GAPDH was detected using moues monoclonal anti-GAPDH #G8795 (diluted 1:3000) and rabbit anti-mouse IgG H&L (HRP) ab6728 (diluted 1:5000). ECL Plus kit was used for the detection.

### Generation of sgRNA constructs

SgRNA constructs targeting *TP53* P2 promoter were generated by cloning the respective gRNA oligonucleotides (gRNA_P2_1F:5’-GTAAGGACTGGGGCGCGGCA-3’; gRNA_P2_1R: 5’-TGCCGCGCCCCAGTCCTTAC-3’; gRNA_P2_2F: 5’-GCGTCTGTTCATTTGACTGC-3’; gRNA_P2_2R: 5’-GCAGTCAAATGAACAGACGC-3’) into pX330-U6-Chimeric_BB-CBh-SpCas9 vector which digested with BbsI from Feng Zhang (Addgene plasmid # 42230; http://n2t.net/addgene:4223O;RRID:Addgene_42230), both sgRNA constructs were cotransfected into pig OS cells.

## Acknowledgments

The authors thank Johanna Tebbing for technical assistance with molecular biology, Steffen and Viola Loebnitz, Gerhard Kammermeier, Konrad Praller and Andres Sohn for animal husbandry.

## Authors’ contributions

KF, AK, DS, J-CB and AS designed the study. GN, IS, TF, BR and AC carried out molecular experiments. KF, GN, KFi and BK collected samples. BS, BB, EK and KS performed histopathological analysis of tumours. RB provided human OS samples. HP performed statistical analysis. KF, TF, AK and AS wrote the manuscript with contributions from other authors. J-CB and DS edited the manuscript. All authors critically reviewed and approved the final manuscript.

## Ethics approval and consent to participate

All experimental procedures were conducted in accordance with the ethical standards, approved by the Government of Upper Bavaria (permit number 55.2-1-54-2532-6-13) and performed according to the German Animal Welfare Act and European Union Normative for Care and Use of Experimental Animals (EU Directive 2010/63/EU).

## Consent for publication

Not applicable

## Legends to supplementary figures

**Figure S1.** Protein sequence alignment of human Δ160p53α and pig Δ152p53α isoform. Dissimilarities between the analysed sequences are indicated as (+) and an empty space.

**Figure S2**. Cross-species alignment of *TP53* intron 4 sequence, at the location of the human P2 promoter. Regions of greatest similarity between human and pig are highlighted in yellow. Predicted transcription factor binding sites are underlined. No binding sites for the listed transcription factors were identified in the mouse intron 4.

**Figure S3.** Silencing of Δ152p53α expression in pig OS cells. (**a**) PCR showing the CRISPR/Cas9 guided deletion of the *TP53* P2 promoter (9665 to 10374bp on the NC_010454) in *flTP53^R167H^* pig OS cells. (**b**) RT-PCR showing the lack of Δ152p53α expression in the edited pig OS cells. For the RT-PCR, primers specific for the Δ152p53 mRNA expression were used. M – marker, ΔP2 - RT-PCR result in osteosarcoma cells with deleted TP53 P2 promoter, P2 – RT-PCR (807bp) fragment of *TP53* in unedited *flTP53^R167H^* pig osteosarcoma cells, nc – negative control.

**Figure S4.** RT-PCR analysis of Δ133/160p53 mRNA expression in human OS samples.

**Figure S5.** Functional analysis of Δ152p53α isoform. Proliferation assay in pig OS cells transfected with an expression vector carrying the wild type Δ152p53α or mutant R167H Δ152p53α cDNA sequence under the control of the CAG promoter. The GFP vector was used as control.

**Figure S6.** DNA methylation analysis of selected genomic *TP53* regions in OS (n= 48) and matched healthy bone samples of *flTP53^R167H/R167H^* pigs. The analysed fragments included CpG sites in promoter 1 **(a),** fragment Pint_1 **(b),** Pint_2 **(c)**, Pint_ 3 **(d)** in intron 1, and exon 5 **(e)**. Values represent mean ± standard deviation. * P < 0.05, ** P < 0.01.

## References

1. Robertson N, Schook LB, Schachtschneider KM. Porcine cancer models: potential tools to enhance cancer drug trials. Expert Opin Drug Discov. 2020;15(8):893–902.

2. Flisikowska T, Kind A, Schnieke A. Pigs as models of human cancers. Theriogenology. 2016;86(1):433–7.

3. Kalla D, Kind A, Schnieke A. Genetically Engineered Pigs to Study Cancer. Int J Mol Sci. 2020;21(2).

4. Rogalla S, Flisikowski K, Gorpas D, Mayer AT, Flisikowska T, Mandella MJ, et al. Biodegradable Fluorescent Nanoparticles for Endoscopic Detection of Colorectal Carcinogenesis. Advanced Functional Materials. 2019;29(51):1904992.

5. Jacks T, Remington L, Williams BO, Schmitt EM, Halachmi S, Bronson RT, et al. Tumor spectrum analysis in p53-mutant mice. Curr Biol. 1994;4(1):1–7.

6. Jackson JG, Lozano G. The mutant p53 mouse as a pre-clinical model. Oncogene. 2013;32(37):4325–30.

7. Hingorani SR, Wang L, Multani AS, Combs C, Deramaudt TB, Hruban RH, et al. Trp53R172H and KrasG12D cooperate to promote chromosomal instability and widely metastatic pancreatic ductal adenocarcinoma in mice. Cancer Cell. 2005;7(5):469–83.

8. Xu X, Qiao W, Linke SP, Cao L, Li WM, Furth PA, et al. Genetic interactions between tumor suppressors Brca1 and p53 in apoptosis, cell cycle and tumorigenesis. Nat Genet. 2001;28(3):266–71.

9. Chen Z, Cheng K, Walton Z, Wang Y, Ebi H, Shimamura T, et al. A murine lung cancer co-clinical trial identifies genetic modifiers of therapeutic response. Nature. 2012;483(7391):613–7.

10. Leuchs S, Saalfrank A, Merkl C, Flisikowska T, Edlinger M, Durkovic M, et al. Inactivation and inducible oncogenic mutation of p53 in gene targeted pigs. PLoS One. 2012;7(10):e43323.

11. Saalfrank A, Janssen KP, Ravon M, Flisikowski K, Eser S, Steiger K, et al. A porcine model of osteosarcoma. Oncogenesis. 2016;5:e210.

12. Donehower LA, Lozano G. 20 years studying p53 functions in genetically engineered mice. Nat Rev Cancer. 2009;9(11):831–41.

13. Olive KP, Tuveson DA, Ruhe ZC, Yin B, Willis NA, Bronson RT, et al. Mutant p53 gain of function in two mouse models of Li-Fraumeni syndrome. Cell. 2004;119(6):847–60.

14. Lang GA, Iwakuma T, Suh YA, Liu G, Rao VA, Parant JM, et al. Gain of function of a p53 hot spot mutation in a mouse model of Li-Fraumeni syndrome. Cell. 2004;119(6):861–72.

15. Guijarro MV, Ghivizzani SC, Gibbs CP. Animal models in osteosarcoma. Front Oncol. 2014;4:189.

16. Mirabello L, Troisi RJ, Savage SA. International osteosarcoma incidence patterns in children and adolescents, middle ages and elderly persons. International journal of cancer Journal international du cancer. 2009;125(1):229–34.

17. Durfee RA, Mohammed M, Luu HH. Review of Osteosarcoma and Current Management. Rheumatol Ther. 2016;3(2):221–43.

18. Bielack SS, Kempf-Bielack B, Delling G, Exner GU, Flege S, Helmke K, et al. Prognostic factors in high-grade osteosarcoma of the extremities or trunk: an analysis of 1,702 patients treated on neoadjuvant cooperative osteosarcoma study group protocols. J Clin Oncol. 2002;20(3):776–90.

19. Li FP, Fraumeni JF, Jr. Soft-tissue sarcomas, breast cancer, and other neoplasms. A familial syndrome? Ann Intern Med. 1969;71(4):747–52.

20. Ognjanovic S, Olivier M, Bergemann TL, Hainaut P. Sarcomas in TP53 germline mutation carriers: a review of the IARC TP53 database. Cancer. 2012;118(5):1387–96.

21. Muller PA, Vousden KH. p53 mutations in cancer. Nat Cell Biol. 2013;15(1):2–8.

22. Bourdon JC, Fernandes K, Murray-Zmijewski F, Liu G, Diot A, Xirodimas DP, et al. p53 isoforms can regulate p53 transcriptional activity. Genes Dev. 2005;19(18):2122–37.

23. Joruiz SM, Bourdon JC. p53 Isoforms: Key Regulators of the Cell Fate Decision. Cold Spring Harb Perspect Med. 2016;6(8).

24. Fujita K, Mondal AM, Horikawa I, Nguyen GH, Kumamoto K, Sohn JJ, et al. p53 isoforms Delta133p53 and p53beta are endogenous regulators of replicative cellular senescence. Nat Cell Biol. 2009;11(9):1135–42.

25. Bernard H, Garmy-Susini B, Ainaoui N, Van Den Berghe L, Peurichard A, Javerzat S, et al. The p53 isoform, Delta133p53alpha, stimulates angiogenesis and tumour progression. Oncogene. 2013;32(17):2150–60.

26. Seva J, Pallares FJ, Gomez MA, Bernabe A. Osteoblastic osteosarcoma in a fattening pig. Vet Rec. 2001;148(5):147–8.

27. Marcel V, Perrier S, Aoubala M, Ageorges S, Groves MJ, Diot A, et al. Delta160p53 is a novel N-terminal p53 isoform encoded by Delta133p53 transcript. FEBS Lett. 2010;584(21):4463–8.

28. Lee DS, Choung HW, Kim HJ, Gronostajski RM, Yang YI, Ryoo HM, et al. NFI-C regulates osteoblast differentiation via control of osterix expression. Stem Cells. 2014;32(9):2467–79.

29. Fane M, Harris L, Smith AG, Piper M. Nuclear factor one transcription factors as epigenetic regulators in cancer. Int J Cancer. 2017;140(12):2634–41.

30. Denny SK, Yang D, Chuang CH, Brady JJ, Lim JS, Gruner BM, et al. Nfib Promotes Metastasis through a Widespread Increase in Chromatin Accessibility. Cell. 2016;166(2):328–42.

31. Poole LA, Cortez D. Functions of SMARCAL1, ZRANB3, and HLTF in maintaining genome stability. Crit Rev Biochem Mol Biol. 2017;52(6):696–714.

32. Seki M, Kimura S, Isobe T, Yoshida K, Ueno H, Nakajima-Takagi Y, et al. Recurrent SPI1 (PU.1) fusions in high-risk pediatric T cell acute lymphoblastic leukemia. Nat Genet. 2017;49(8):1274–81.

33. Peng Y, Chen L, Li C, Lu W, Agrawal S, Chen J. Stabilization of the MDM2 oncoprotein by mutant p53. J Biol Chem. 2001;276(9):6874–8.

34. Blackburn J, Roden DL, Ng R, Wu J, Bosman A, Epstein RJ. Damage-inducible intragenic demethylation of the human TP53 tumor suppressor gene is associated with transcription from an alternative intronic promoter. Mol Carcinog. 2016;55(12):1940–51.

35. Tornaletti S, Pfeifer GP. Complete and tissue-independent methylation of CpG sites in the p53 gene: implications for mutations in human cancers. Oncogene. 1995;10(8):1493–9.

36. Zhang J, Liu H, Hou L, Wang G, Zhang R, Huang Y, et al. Circular RNA_LARP4 inhibits cell proliferation and invasion of gastric cancer by sponging miR-424-5p and regulating LATS1 expression. Mol Cancer. 2017;16(1):151.

37. Chen N, Zhao G, Yan X, Lv Z, Yin H, Zhang S, et al. A novel FLI1 exonic circular RNA promotes metastasis in breast cancer by coordinately regulating TET1 and DNMT1. Genome Biol. 2018;19(1):218.

38. Chen S, Moroi Y, Urabe K, Takeuchi S, Kido M, Hayashida S, et al. Differential expression of two new members of the p53 family, p63 and p73, in extramammary Paget’s disease. Clin Exp Dermatol. 2008;33(5):634–40.

39. Jost CA, Marin MC, Kaelin WG, Jr. p73 is a simian [correction of human] p53-related protein that can induce apoptosis. Nature. 1997;389(6647):191–4.

40. Flores ER, Sengupta S, Miller JB, Newman JJ, Bronson R, Crowley D, et al. Tumor predisposition in mice mutant for p63 and p73: evidence for broader tumor suppressor functions for the p53 family. Cancer Cell. 2005;7(4):363–73.

41. Irwin MS. DeltaNp73: misunderstood protein? Cancer Biol Ther. 2006;5(7):804–7.

42. Conforti F, Yang AL, Agostini M, Rufini A, Tucci P, Nicklison-Chirou MV, et al. Relative expression of TAp73 and DeltaNp73 isoforms. Aging (Albany NY). 2012;4(3):202–5.

43. Moll UM. The Role of p63 and p73 in tumor formation and progression: coming of age toward clinical usefulness. Commentary re: F. Koga et al., Impaired p63 expression associates with poor prognosis and uroplakin III expression in invasive urothelial carcinoma of the bladder. Clin. Cancer Res., 9: 5501-5507, 2003, and P. Puig et al., p73 Expression in human normal and tumor tissues: loss of p73alpha expression is associated with tumor progression in bladder Cancer. Clin. Cancer Res., 9: 5642-5651, 2003. Clin Cancer Res. 2003;9(15):5437–41.

44. Slatter TL, Hung N, Bowie S, Campbell H, Rubio C, Speidel D, et al. Delta122p53, a mouse model of Delta133p53alpha, enhances the tumor-suppressor activities of an attenuated p53 mutant. Cell Death Dis. 2015;6:e1783.

45. Slatter TL, Hung N, Campbell H, Rubio C, Mehta R, Renshaw P, et al. Hyperproliferation, cancer, and inflammation in mice expressing a Delta133p53-like isoform. Blood. 2011;117(19):5166–77.

46. Campbell HG, Slatter TL, Jeffs A, Mehta R, Rubio C, Baird M, et al. Does Delta133p53 isoform trigger inflammation and autoimmunity? Cell Cycle. 2012;11(3):446–50.

47. Bykov VJN, Eriksson SE, Bianchi J, Wiman KG. Targeting mutant p53 for efficient cancer therapy. Nat Rev Cancer. 2018;18(2):89–102.

48. Anbarasan T, Bourdon JC. The Emerging Landscape of p53 Isoforms in Physiology, Cancer and Degenerative Diseases. Int J Mol Sci. 2019;20(24).

49. Fragou A, Tzimagiorgis G, Karageorgopoulos C, Barbetakis N, Lazopoulos A, Papaioannou M, et al. Increased Delta133p53 mRNA in lung carcinoma corresponds with reduction of p21 expression. Mol Med Rep. 2017;15(4):1455–60.

50. Hofstetter G, Berger A, Schuster E, Wolf A, Hager G, Vergote I, et al. Delta133p53 is an independent prognostic marker in p53 mutant advanced serous ovarian cancer. Br J Cancer. 2011;105(10):1593–9.

51. Nutthasirikul N, Limpaiboon T, Leelayuwat C, Patrakitkomjorn S, Jearanaikoon P. Ratio disruption of the 133p53 and TAp53 isoform equilibrium correlates with poor clinical outcome in intrahepatic cholangiocarcinoma. Int J Oncol. 2013;42(4):1181–8.

52. Avery-Kiejda KA, Morten B, Wong-Brown MW, Mathe A, Scott RJ. The relative mRNA expression of p53 isoforms in breast cancer is associated with clinical features and outcome. Carcinogenesis. 2014;35(3):586–96.

53. Candeias MM, Hagiwara M, Matsuda M. Cancer-specific mutations in p53 induce the translation of Delta160p53 promoting tumorigenesis. EMBO Rep. 2016;17(11):1542–51.

54. Sieren JC, Meyerholz DK, Wang XJ, Davis BT, Newell JD, Jr., Hammond E, et al. Development and translational imaging of a TP53 porcine tumorigenesis model. J Clin Invest. 2014;124(9):4052–66.

55. Stindt MH, Muller PA, Ludwig RL, Kehrloesser S, Dotsch V, Vousden KH. Functional interplay between MDM2, p63/p73 and mutant p53. Oncogene. 2015;34(33):4300–10.

56. Zhang J, Sun W, Kong X, Zhang Y, Yang HJ, Ren C, et al. Mutant p53 antagonizes p63/p73-mediated tumor suppression via Notch1. Proc Natl Acad Sci U S A. 2019;116(48):24259–67.

57. Gonfloni S, Caputo V, Iannizzotto V. P63 in health and cancer. Int J Dev Biol. 2015;59(1-3):87–93.

58. Lucena-Araujo AR, Kim HT, Thome C, Jacomo RH, Melo RA, Bittencourt R, et al. High DeltaNp73/TAp73 ratio is associated with poor prognosis in acute promyelocytic leukemia. Blood. 2015;126(20):2302–6.

59. Robinson D, Van Allen EM, Wu YM, Schultz N, Lonigro RJ, Mosquera JM, et al. Integrative clinical genomics of advanced prostate cancer. Cell. 2015;161(5):1215–28.

60. Amin MB, Greene FL, Edge SB, Compton CC, Gershenwald JE, Brookland RK, et al. The Eighth Edition AJCC Cancer Staging Manual: Continuing to build a bridge from a population-based to a more “personalized” approach to cancer staging. CA Cancer J Clin. 2017;67(2):93–9.

