## Supplemental Information for "Porcine model elucidates function of p53 isoform in carcinogenesis"

```
              Pos.
        Human 160   MAIYKQSQHMTEVVRRCPHHERCSD-SDGLAPPQHLIRVEGNLRVEYLDDRNTFRHSVVV
                    MAIYK+S++MTEVVRRCPHHER SD SDGLAPPQHLIRVEGNLR EYLDDRNTFRHSVVV
        Pig   152   MAIYKKSEYMTEVVRRCPHHERSSDYSDGLAPPQHLIRVEGNLRAEYLDDRNTFRHSVVV

        Human 219   PYEPPEVGSDCTTIHYNYMCNSSCMGGMNRRPILTIITLEDSSGNLLGRNSFEVRVCACP
                    PYEPPEVGSDCTTIHYN+MCNSSCMGGMNRRPILTIITLED+SGNLLGRNSFEVRVCACP
        Pig   212   PYEPPEVGSDCTTIHYNFMCNSSCMGGMNRRPILTIITLEDASGNLLGRNSFEVRVCACP

        Human 279   GRDRRTEEENLRKKGEPHHELPPGSTKRALPNNTSSSPQPKKKPLDGEYFTLQIRGRERF
                    GRDRRTEEEN  KKG+   E PPGSTKRALP +TSSSP  KKKPLDGEYFTLQIRGRERF
        Pig   272   GRDRRTEEENFLKKGQSCPEPPPGSTKRALPTSTSSSPVQKKKPLDGEYFTLQIRGRERF

        Human 339   EMFRELNEALELKDAQAGKEPGGSRAHSSHLKSKKGQSTSRHKKLMFKTEGPDSD  393
                    EMFRELN+ALELKDAQ  +E G +RAHSSHLKSKKGQS SRHKK MFK EGPDSD
        Pig   332   EMFRELNDALELKDAQTARESGENRAHSSHLKSKKGQSPSRHKKPMFKREGPDSD  386
```

**Figure S1.** Protein sequence alignment of human Δ160p53α and pig Δ152p53α isoform. Dissimilarities between the analysed sequences are indicated as (+) and an empty space.

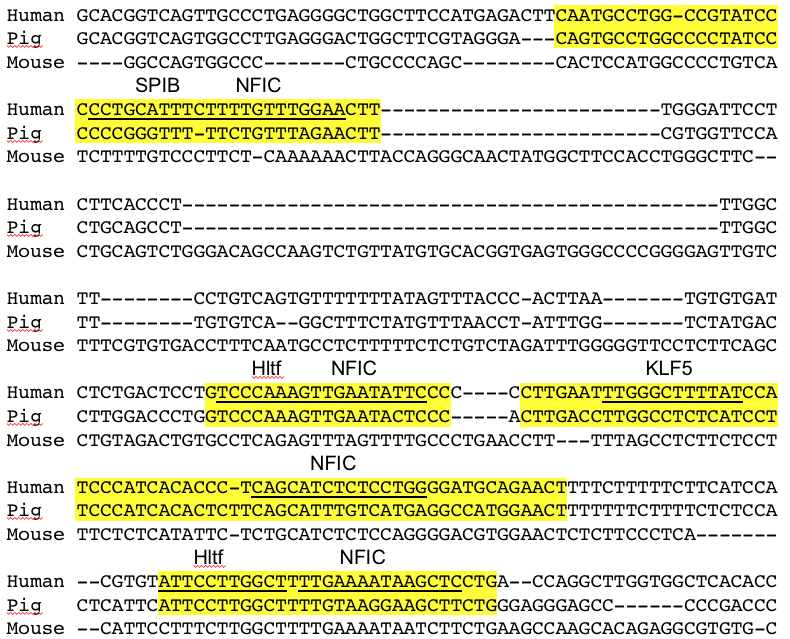

**Figure S2**. Cross-species alignment of *TP53* intron 4 sequence, at the location of the human P2 promoter. Regions of greatest similarity between human and pig are highlighted in yellow. Predicted transcription factor binding sites are underlined. No binding sites for the listed transcription factors were identified in the mouse intron 4.

a

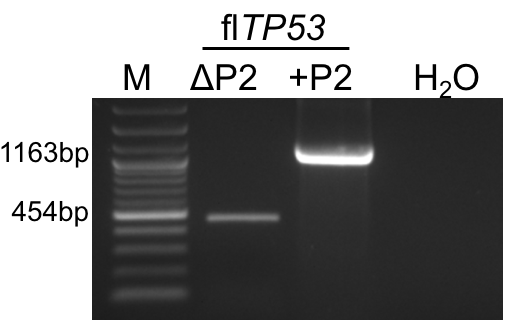

b

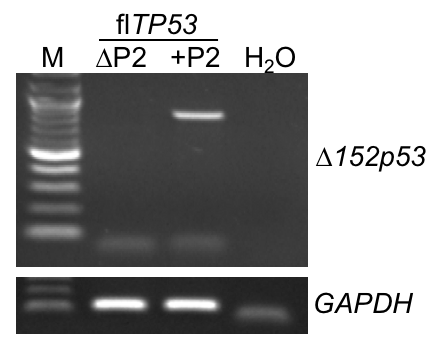

**Figure S3.** Silencing of Δ152p53α expression in pig OS cells. (**a**) PCR showing the CRISPR/Cas9 guided deletion of the *TP53* P2 promoter (9665 to 10374bp on the NC_010454) in *flTP53^R167H^* pig OS cells. (**b**) RT-PCR showing the lack of Δ152p53α expression in the edited pig OS cells. For the RT-PCR, primers specific for the Δ152p53 mRNA expression were used. M - marker, ΔP2 - RT-PCR result in osteosarcoma cells with deleted TP53 P2 promoter, P2 – RT-PCR (807bp) fragment of *TP53* in unedited *flTP53^R167H^* pig osteosarcoma cells, nc – negative control.

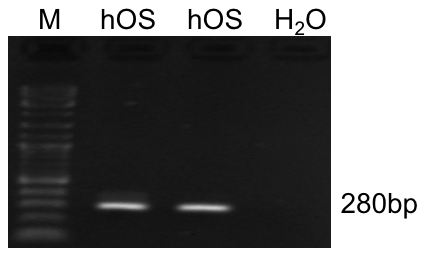

**Figure S4**. RT-PCR analysis of Δ133/160p53 mRNA expression in human OS samples.

*

**

Cell number (x 105)

|  | GFP | ∆152TP53WT | ∆152TP53R167H |
| --- | --- | --- | --- |
| 0h | 0.5 | 0.5 | 0.5 |
| 24h | 0.7 | 0.633333 | 0.9 |
| 48h | 1.1 | 1 | 1.06667 |
| 72h | 1.9 | 1.86667 | 2.23333 |
| 96h | 3.83333 | 4.93333 | 4.73333 |
| 120h | 5.03333 | 6.53333 | 7.93333 |

**Figure S5**. Functional analysis of Δ152p53α isoform. Proliferation assay in pig OS cells transfected with an expression vector carrying the wild type Δ152p53α or mutant R167H Δ152p53α cDNA sequence under the control of the CAG promoter. The GFP vector was used as control.

a

DNA methylation (%)

|  | bone | osteosarcoma |
| --- | --- | --- |
| CpG1 | 11 | 14 |
| CpG2 | 18 | 17 |
| CpG3 | 9 | 24 |
| CpG4 | 15 | 21 |
| CpG5 | 18 | 18 |
| CpG6 | 22 | 26 |

b

*

DNA methylation (%)

|  | Healthy bone | Osteosarcoma |
| --- | --- | --- |
| CpG1 | 45 | 31 |
| CpG2 | 30 | 20 |
| CpG3 | 43 | 29 |
| CpG4 | 24 | 16 |
| CpG5 | 67 | 61 |
| CpG6 | 28 | 24 |

c

DNA methylation (%)

|  | Healthy bone | Osteosarcoma |
| --- | --- | --- |
| CpG1 | 77 | 75 |
| CpG2 | 89 | 85 |
| CpG3 | 91 | 93 |
| CpG4 | 88 | 85 |
| CpG5 | 75 | 83 |

d

DNA methylation (%)

|  | Healthy bone | Osteosarcoma |
| --- | --- | --- |
| CpG1 | 95 | 93 |
| CpG2 | 92 | 96 |
| CpG3 | 96 | 98 |

e

**

**

DNA methylation (%)

|  |  |  |
| --- | --- | --- |
| CpG1 | 91 | 84 |
| CpG2 | 92 | 88 |
| CpG3 | 99 | 93 |
| CpG4 | 92 | 87 |
| CpG5 | 96 | 63 |
| CpG6 | 69 | 48 |

**Figure S6**. DNA methylation analysis of selected genomic *TP53* regions in OS (n= 48) and matched healthy bone samples of *flTP53^R167H/R167H^* pigs. The analysed fragments included CpG sites in promoter 1 **(a),** fragment Pint_1 **(b),** Pint_2 **(c)**, Pint_ 3 **(d)** in intron 1, and exon 5 **(e)**. Values represent mean ± standard deviation. * P < 0.05, ** P < 0.01.
